# Equilibration-free cryopreservation of beef and bison semen

**DOI:** 10.64898/2026.05.15.725595

**Authors:** S Yang, K Rajapaksha, E Zwiefelhofer, GP Adams, M Anzar

## Abstract

Conventional semen cryopreservation involves equilibration at 4°C and optimum freezing rates. We hypothesized that a cholesterol-based semen extender obviates the need for equilibration, minimizing total processing time for semen cryopreservation. Experiments were conducted to determine the effects of semen extender (egg yolk- or cholesterol-based) and freezing method (routine or fast) on post-thaw sperm characteristics and fertility of beef and bison semen. In Experiment 1, beef semen diluted in tris-egg yolk-glycerol (TEYG) or cholesterol-cyclodextrin tris-glycerol (CCTG) extender underwent routine or fast freezing method. Cholesterol from animal and plant origins were compared. The routine method included 90-min equilibration at 4°C and routine freezing (RE-RF, total time 97 min) whereas the fast method included no equilibration and fast freezing (NE-FF, total time 14 min). Post-thaw sperm quality was assessed by CASA, and *in vitro* fertilization. Post-thaw sperm motility was not affected by the origin of cholesterol (animal or plant), but was lowest in the TEYG NE-FF group (24% vs 43-51%, P < 0.05). *In vitro* cleavage and blastocyst development rates did not differ between RE-RF and NE-FF groups. In Experiment 2, bison semen was diluted in TEYG or plant-CCTG extender and frozen as in Experiment 1. Post-thaw sperm motility was lowest in the TEYG NE-FF group (10% vs 39-51%, P < 0.05). In Experiment 3, beef semen diluted in TEYG or plant-CCTG extender underwent either a routine (RE-RF) or modified freezing (NE-RF, total time 25 min) method. Post-thaw sperm characteristics did not differ between extenders but were greater using routine freezing (RE-RF) compared to the modified method of freezing (NE-RF). Pregnancy rates were similar between extenders (TEYG vs plant-CCTG) using the modified freezing method without equilibration and insemination at 72 h after progesterone device removal. In conclusion, beef and bison semen diluted in cholesterol-based extender may be cryopreserved without equilibration.

## Introduction

Mammalian semen cryopreservation procedures include dilution in cryoprotective extenders, cooling to 4°C, and freezing below 0°C. Semen extenders increase both ejaculate volume and sperm longevity. During initial cooling to 4°C, sperm plasma membranes biochemically interact with extender constituents, specifically referred to as ‘equilibration’ [1]. Mammalian sperm plasma membranes undergo lipid-phase transitions between 18° to 14°C, known as ‘cold shock’ [2–4], characterized by lateral movement of phospholipids which increase membrane permeability to ions, and damages sperm membranes irreversibly [5,6]. Upon ejaculation, binder of sperm proteins in seminal plasma causes efflux of phospholipids and cholesterol from plasma membranes. Egg yolk in extender mitigates the cold shock effect by a mechanism that involves sequestering binder of sperm proteins in seminal plasma [7,8] and binding with sperm plasma membranes [9], thus stabilizing sperm against membrane lipid-phase transitions and replacing phospholipids lost during freezing and thawing [10–13].

After dilution in extender, semen is commonly held at 4°C for 90-120 min to achieve equilibrium between components in semen extender and sperm plasma membranes. With egg yolk-glycerol extenders, the equilibration time is commonly ≥90 min before freezing below 0°C [14]. However, a wide range of equilibration time has been reported in literature; long equilibration time (>12 h) provided better post-thaw sperm quality than short time [1,15–19]. In a typical semen cryopreservation protocol, the prerequisite equilibration step prolongs the total processing time and negatively impacts the efficiency of commercial semen production centers.

Earlier work demonstrated that exogenous cholesterol is incorporated into sperm plasma membranes within 15 min of incubation in the presence of egg yolk components, and improved post-thaw sperm motility in beef and bison bulls [20–22]. However, the use of egg yolk, an animal product, in semen extender raises biosecurity concerns regarding transmission of infectious diseases. Moreover, egg yolk has undefined composition and varies from batch to batch. Cholesterol, commonly from sheep’s wool, is known to modulate membrane fluidity, increase membrane stability, and minimize lipid-phase transitions during cooling [23]. Alternatively, cholesterol may be sourced from plants to minimize biosecurity risks associated with animal products and have a defined composition. Cyclodextrins act as carriers to deliver cholesterol to sperm plasma membrane [22–24]. In this regard, we developed a novel cholesterol-based extender as an alternative to conventional egg yolk extender for cryopreservation of beef and bison semen to mitigate biosecurity risks [11,25].

After equilibration, sperm survival below 0°C depends primarily on minimizing two physico-chemical effects: intracellular ice crystal formation and high solute concentrations [26]. If cell freezing occurs at an overly fast rate, incomplete dehydration leads to intracellular ice formation of residual water, causing physical damage to the cell membrane and organelles. Conversely, if the freezing rate is too slow, prolonged intracellular dehydration leads to high solute concentrations (solution effect) and toxicity. Glycerol, a low-molecular weight penetrating cryoprotectant, readily crosses the plasma membrane, binds with free water and helps in preventing ice crystal formation, and can be added either before or after cooling [1,25,27,28]. Therefore, total processing time may be further reduced by implementing a fast freezing rate below 0°C.

North American wood and plains bison are threatened by genetic isolation and disease, and concerted efforts are underway to rescue the species by establishing a bison biobank with the use of germplasm in assisted reproductive technologies [25,29–31]. However, semen collection from free-roaming bison under wild conditions poses logistical concerns related to cryopreservation processing time and facilities. There is a need for a simple and quick method of semen cryopreservation to enable collection and processing under remote field conditions and during extremely cold months.

In the present study, attempts were made to develop a short cryopreservation protocol for bovine and bison semen by removing the most time-consuming equilibration step, and by freezing at a faster rate. We hypothesized that a cholesterol-based semen extender obviates the need for equilibration minimizing total processing time for semen cryopreservation. The specific objectives of this study were to determine the effects of semen extender (egg yolk- or cholesterol-based), origin of cholesterol (animal or plant), and freezing method (routine or fast) on post-thaw sperm quality and fertility of beef and bison semen.

## Materials and Methods

Beef cattle and bison bulls were housed separately (5 km apart) under similar management and nutrition conditions at the Livestock and Forage Center of Excellence, University of Saskatchewan, Saskatoon. Animal procedures were conducted in accordance with the Canadian Council on Animal Care and approved by the University of Saskatchewan Animal Care Committee (Animal Use Protocol #20100150). In all experiments, semen was collected by electroejaculation (Pulsator IV Auto Adjust; Lane Manufacturing Inc., Denver, CO, USA). After collection, semen was transported to the Cryobiology Laboratory, Westgen Research Suite, University of Saskatchewan, within 2 h at 32°C. Ejaculates were evaluated by computer-assisted sperm analyzer (CASA; Sperm Vision 3.0, Minitube Canada, Ingersoll, ON, Canada), as reported earlier [20,32]. Ejaculates possessing sperm concentration >200×10^6^ sperm/mL and total motility >60% were pooled to minimize bull-to-bull differences, for further processing. All chemicals were purchased from Sigma-Aldrich (Oakville, ON, Canada) unless otherwise stated.

### Semen extender preparation

Cholesterol-cyclodextrin complex (CC) was prepared as previously described [21]. Solution A was prepared by dissolving 200 mg cholesterol of animal-origin (sheep wool; Cat# C8667) or plant-origin (PhytoChol^®^, Wilshire Technologies, Princeton, NJ, USA; Cat# 57-88-5) in 1 mL chloroform. Solution B was prepared by dissolving 1 g methyl β-cyclodextrin (Cat# C4555) in 2 mL methanol. Solution A (0.45 mL) was then added to Solution B (2 mL) and mixed until the solution became homogenous. The mixture was then poured into a glass petri dish and dried under a gentle stream of nitrogen gas. The resulting crystals were desiccated overnight and stored in a glass bottle in desiccator at 22°C until used. On the day of experiment, a working solution of CC (50 mg/mL) was prepared in tris-citric acid (TCA) buffer containing tris-base 3.03% wt/vol, citric acid monohydrate 1.74% wt/vol, and fructose 1.2% wt/vol (pH 7.1) in Milli-Q distilled water and used immediately.

Tris-glycerol (TG; 2×) extender was prepared by adding glycerol (14% v/v), gentamycin sulfate (1 mg/mL), tylosin (200 µg/mL; Tylan Soluble, Elanco, Guelph, ON, Canada), and lincomycin-spectinomycin (600/1200 µg/mL; Linco-Spectin, Pfizer Animal Health, Kirkland, QC, Canada) to TCA buffer, and stored at -20°C. TG (2×) extender was added to CC treated semen (1:1) to achieve the final concentrations of glycerol (7% v/v), gentamycin sulfate (500 µg/mL), tylosin (100 µg/mL), and lincomycin-spectinomycin (300/600 µg/mL). The combined extender will be referred to as ‘CCTG’ hereafter.

A conventional tris-egg yolk-glycerol (TEYG) extender was prepared by adding glycerol (7%, v/v), egg yolk (20%, v/v), gentamycin sulfate (500 µg/mL), tylosin (100 µg/mL), and lincomycin-spectinomycin (300/600 µg/mL) in TCA buffer. The final extender was centrifuged at 12,000× *g* for 15 min at 4°C. The supernatant was recovered and stored at -20°C. All frozen media were thawed to 37°C on the day of the experiment.

### Experiment 1: Effect of extender, origin of cholesterol and freezing method on post-thaw sperm quality and *in vitro* fertility of beef bulls

#### Experiment 1a (Simmental bulls)

Semen was collected from 5 Simmental bulls on 5 different dates (replicates) to determine the effects of semen extender (egg yolk- or cholesterol-based), source of cholesterol (animal or plant) and freezing method (routine or fast) on post-thaw sperm motility, and plasma membrane and acrosome integrity. Pooled ejaculates were divided into three extender groups: TEYG, animal-CCTG, or plant-CCTG, as previously described [11,25]. For the TEYG group, semen was diluted to 50×10^6^ sperm/mL with extender (32°C) and held at 22°C for 15 min. For animal-and plant-CCTG groups, semen was initially diluted to 100×10^6^ sperm/mL with TCA buffer at 32°C, treated with 1 mg animal- or plant-CCTG/ml semen at 22°C for 15 min, and then diluted 1:1 with TG (2×) to achieve a final sperm concentration of 50×10^6^/mL [25]. The final concentration of animal- or plant-CC was 0.5 mg/ml semen.

Each TEYG- and CCTG-diluted batch of semen underwent either a routine (routine equilibration-routine freezing; RE-RF) or fast freezing (no equilibration-fast freezing; NE-FE) method (Table 1). In the routine method, 15-ml tubes of extended semen were placed in a 500 mL glass beaker containing water (22°C) and cooled at 4°C for 90 min (equilibration), then filled into 0.5 mL straws and frozen using a routine freezing curve (i.e., -3°C/min from 4°C to -10°C, - 40°C/min from -10°C to -80°C) in a programmable cell freezer (ICE-CUBE 14-S; Sy-lab Version 1.30, Gerate GmbH, Neupurkdersdof, Austria) [11]. In the fast method, diluted semen was immediately filled into 0.5 mL straws at 22°C, and frozen directly in a programmable cell freezer without equilibration at 4°C (i.e., -1°C/min from 22°C to 10°C, and -40°C/min from 10°C to -80°C). After reaching -80°C in both freezing methods, semen straws were plunged into liquid nitrogen, and stored until thawing.

**Table 1.** Duration of routine and fast freezing methods for cryopreservation of beef and bison semen (Experiments 1 & 2).

| Stage of cryopreservation | Routine method<br>(RE-RF)* | Fast method<br>(NE-FF)* |
| --- | --- | --- |
| Cooling/equilibration<br>(time) | 22°C to 4°C<br>(90 min) | 22°C to 10°C @ -1°C/min<br>(12 min) |
| Freezing ramp 1<br>(time) | 4°C to -10°C @ -3°C/min<br>(~5 min) | 10°C to -80°C @ -40°C/min<br>(~2 min) |
| Freezing ramp 2<br>(time) | -10°C to -80°C @ -40°C/min<br>(~2 min) | - |
| Total processing time | ~97 min | ~14 min |

\*RE-RF: routine equilibration-routine freezing; NE-FF: no equilibration-fast freezing.

For post-thaw sperm analysis, two straws from each treatment group were thawed at 37°C for 1 min and contents were pooled to minimize straw-to-straw variation. Sperm motility was determined at 0 and 2 h post-thaw by CASA using bull settings [16,32], and *in vitro* fertilization potential of sperm in all treatment groups was determined [25,33,34]. Both CASA and IVF procedures are described under the section below (*Semen assays*).

#### Experiment 1b (Angus bulls)

Semen was collected from 5 Angus bulls on 7 different dates (replicates) and pooled, as described above. Since the effect of CC origin (animal- or plant-derived) was not significant for any endpoint in Experiment 1a, subsequent experiments were conducted using plant-CCTG extender for enhanced biosecurity. The effects of semen extender (egg yolk or plant-CCTG) and freezing method (routine or fast) on post-thaw sperm motility and structural characteristics (intact plasma and acrosome membranes) were evaluated at 0 and 2 h, by CASA and flow cytometer respectively, as described below (*Semen assays*).

### Experiment 2: Effect of extender and freezing method on post-thaw sperm quality of bison bulls

Semen was collected from 5 bison bulls on 5 different dates and pooled, as described above. Pooled semen was diluted in either TEYG or plant-CCTG extender and underwent routine or fast method (Table 1). As in Experiment 1, post-thaw sperm motility, and plasma membrane and normal acrosome were assessed by CASA and flow cytometry respectively, at 0 and 2 h, as described below (*Semen assays*).

### Experiment 3: Effect of extender and modified freezing method on post-thaw sperm quality and *in vivo* fertility of beef bulls

Semen was collected from 6 Angus bulls on 10 different dates (replicates) and pooled, as described above. Pooled semen was diluted in either TEYG or plant-CCTG extender and frozen using either a routine (RE-RF) or modified (no equilibration-routine freezing, NE-RF) method (Table 2). Semen straws were deep-frozen using a routine freezing curve, as described in Experiment 1. In the modified group, the extended semen was loaded immediately into 0.5 mL straws at 22°C and cooled from 22°C to 4°C @ -1°C/min using a programmable cell freezer (without equilibration). Below 0°C, the routine freezing curve was used, as described above. Post-thaw sperm motility, and plasma membrane integrity and normal acrosomes were determined using CASA and flow cytometry respectively, at 0 and 2 h, as described below (*Semen assays*). This experiment was repeated on ten pooled ejaculates (replicates) from four Angus bulls, on different dates.

**Table 2.** Duration of routine and modified freezing methods of for cryopreservation of beef semen (Experiment 3).

| Stage of cryopreservation | Routine method<br>(RE-RF)* | Modified method<br>(NE-RF)* |
| --- | --- | --- |
| Cooling<br>(time) | 22°C to 4°C<br>(90 min) | 22°C to 4°C @ -1°C/min<br>(18 min) |
| Freezing ramp 1<br>(time) | 4°C to -10°C @ -3°C/min<br>(~5 min) | 4°C to -10°C @ -3°C/min<br>(~5 min) |
| Freezing ramp 2<br>(time) | -10°C to -80°C @ -40°C/min<br>(~2 min) | -10°C to -80°C @ -40°C/min<br>(~2 min) |
| Total processing time | ~97 min | ~25 min |

\*RE-RF: routine equilibration-routine freezing; NE-RF: no equilibration-routine freezing.

#### Fertility trial 1

A fertility trial was conducted to compare the effect of routine semen processing (TEYG extender and routine freezing [RE-RF]) vs modified semen processing (plant-CCTG extender, modified freezing [NE-RF]) on pregnancy rate after fixed-time insemination. In addition, two different intravaginal progesterone devices (CIDR [Zoetis Canada Inc., Kirkland, QC, Canada] or PRID-Delta [Ceva Animal Health, Cambridge, ON, Canada]) were compared as part of a separate study. Lactating Angus-cross cows (n = 206) at random stages of the estrous cycle and at least 40 days post-partum were divided into 3 replicates and synchronized using a standard 5-day progesterone-based protocol with either a CIDR (n = 105) or PRID-Delta (n = 101; Fig. 1). On Day 0, cows were given GnRH (100 µg gonadorelin *im,* Fertilin) and an intravaginal progesterone device. The progesterone device was removed on Day 5, and cows were given a luteolytic dose of PGF2α (500 µg cloprostenol *im*, Bioestrovet; Vetoquinol, Lavaltrie, QC, Canada) at the time of device removal and again 24 h later. On Day 8 (72 h after PGF2α treatment), cows were treated with GnRH (100 µg gonadorelin *im,* Fertilin) and inseminated with a single dose of TEYG routine (n = 104) or plant-CCTG modified (n = 102) semen, randomly distributed between synchronization treatments. Ovulation was confirmed by transrectal ultrasonography at the time of insemination or subsequent examinations 24 and 48 h post-insemination (MyLab Five, Esaote North America Inc, Fishers, IN, USA), defined as the disappearance of a large follicle between successive examinations or detection of a new CL. Pregnancy was diagnosed by transrectal ultrasonography at 27 to 33 days after insemination. The pregnancy rate was calculated as number pregnant out of those inseminated.

**Fig 1.**
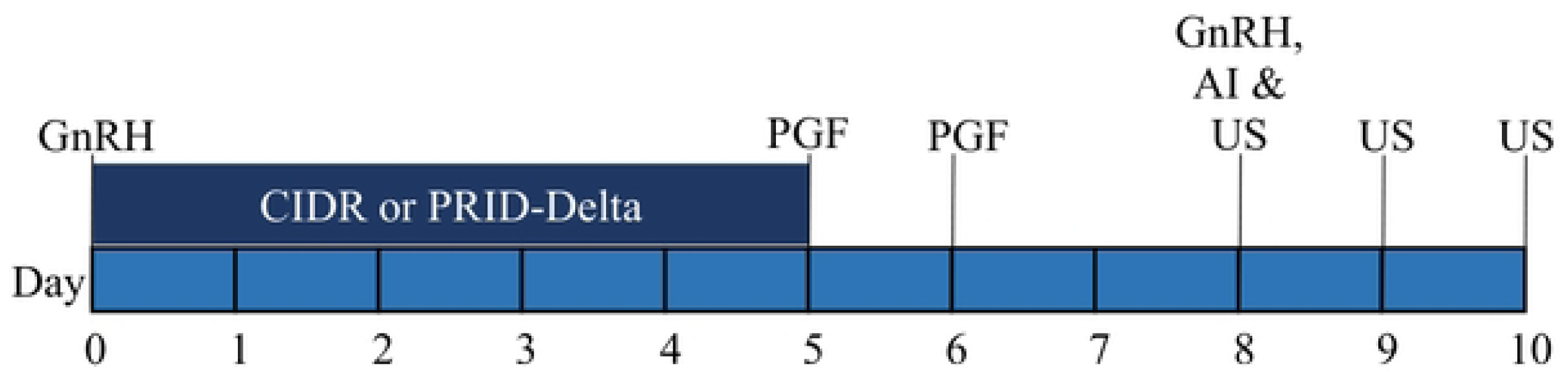
Fixed-time artificial insemination synchronization protocol. Cows were treated with GnRH at the time of placement of a progesterone-releasing intravaginal device (CIDR, n = 105 or PRID-Delta, n = 101). A luteolytic dose of PGF2α was given at the time of device removal and again 24 h later. Cows were treated with GnRH at the time of insemination 72 h after device removal using semen processed in 2 different ways (TEYG routine [RE-RF], n = 104 or plant-CCTG modified [NE-RF], n = 102; Fertility trial 1). Abbreviations: AI, Artificial insemination; CIDR, Controlled internal drug release; GnRH, Gonadotropin-releasing hormone; PGF, Prostaglandin F_2_α; PRID, Progesterone-releasing intravaginal device; US, Ultrasound.

#### Fertility trial 2

A follow-up 2×2 fertility trial was conducted to determine the effect of modified semen (no equilibration-routine freezing, NE-RF) extended in either TEYG or plant-CCTG extenders, and insemination time (60 h or 72 h after prostaglandin treatment) on pregnancy rate after fixed-time artificial insemination. Lactating multiparous Hereford-cross cows (n = 106) at random stages of the estrous cycle and at least 45 days post-partum, were synchronized (Fig. 2). A progesterone-releasing intravaginal device (CIDR, Vetoquinol, Lavaltrie, QC, Canada) was inserted on Day 0 and removed on Day 5. A luteolytic dose of PGF2α (500 µg cloprostenol *im*, Estroplan, Vetoquinol, Lavaltrie, QC, Canada) was given at the time of device removal. Cows were then assigned randomly to be inseminated at either 60 h (n = 65) or 72 h (n = 41) after PGF2α treatment with either TEYG modified (n = 52) or plant-CCTG modified (n = 54) semen group, and treated with GnRH (100 µg gonadorelin *im,* Fertilin, Vetoquinol, Lavaltrie, QC, Canada) at the time of insemination. Ovulation status and pregnancy rates were determined as described in Fertility trial 1.

**Fig 2.**
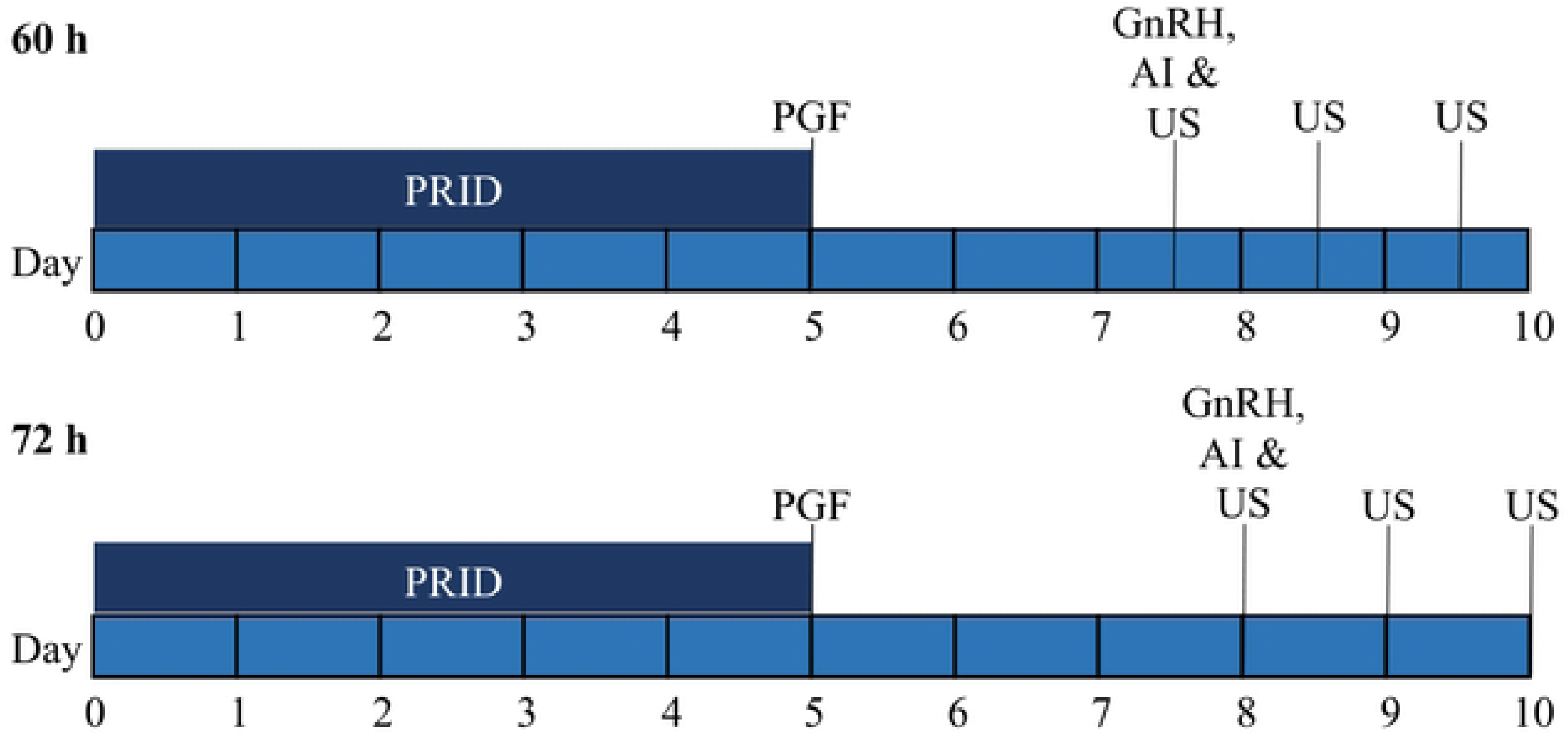
Fixed-time artificial insemination synchronization protocol. A progesterone-releasing intravaginal device (PRID) was placed for a period of 5 days and a luteolytic dose of PGF2α was given at the time of device removal. Cows were treated with GnRH at the time of insemination at either 60 h (n = 65) or 72 h (n = 41) after device removal, using modified (NE-RF) semen diluted TEYG (n = 52) or plant-CCTG (n = 54) extenders (Fertility trial 2). Abbreviations: AI, Artificial insemination; GnRH, Gonadotropin-releasing hormone; PGF, Prostaglandin F_2_α; PRID, Progesterone-releasing intravaginal device; US, Ultrasound.

### Semen assays

#### Computer-assisted sperm analysis

In all experiments, a semen sample (2.5 µl) was placed on a prewarmed (37°C) Leja chamber slide (depth 20 µm; Leja Products B.V. Nieuw-Vennep, Netherlands) and analyzed using a computer-assisted sperm analyzer (CASA; Sperm Vision1 3.0, Minitube Canada). A minimum 200 sperm in 7 fields were assessed for total motility (%; all moving sperm) and progressive motility (%; sperm moving in straight-line, i.e., more than 10 µm radius with a speed of > 4.5 µm/s).

#### Flow cytometer analysis

In Experiments 1b, 2 and 3, plasma and acrosome membrane integrity were assessed by flow cytometer (CyFlow Space, Partec GmbH, Münster, Germany) using propidium iodide (PI; stock 2.4 mM in water) and fluorescein isothiocyanate-peanut agglutinin (FITC-PNA; stock 1 mg/mL in PBS) fluorescent markers, as previously described [16]. Briefly, 1×10^6^ sperm from each treatment group were diluted in 500 µL TCA buffer and incubated with 5 µL PI and 1 µL FITC-PNA at 22°C in the dark for 20 min. Sperm were fixed by adding 10 µL 10% formaldehyde (v/v). Data acquired by FloMax software (version 2.4, Partec GmbH, Münster, Germany) revealed four different sperm populations based on plasma and acrosome membrane integrities: i) sperm with intact plasma membrane and intact acrosome (PI^-^/FITC-PNA^-^, intact), ii) sperm with intact plasma membrane and reacted acrosomes (PI^-^/FITC^-^PNA^+^, partially intact), iii) sperm with compromised plasma membrane and intact acrosomes (PI^+^/FITC-PNA^-^, partially intact), and iv) sperm with compromised plasma membrane and reacted acrosome (PI^+^/FITC-PNA^+^; compromised). Sperm population with an intact plasma membrane and intact acrosome (PI^-^/FITC-PNA^-^; intact) was selected for further statistical analysis.

In all experiments, a stress test was conducted by holding frozen-thawed semen at 37°C for 2 h and analyzed for sperm motility and/or structural characteristics with CASA and flow cytometer respectively.

### *In vitro* fertilization

In Experiment 1a, *in vitro* fertilization potential of frozen-thawed semen extended in TEYG, animal-CCTG, or plant-CCTG extender, and frozen with either routine or fast method were determined. Cattle ovaries were collected from a slaughterhouse near Calgary, Alberta and transported by air to the Cryobiology Laboratory within 8 h, at 22°C. *In vitro* maturation, fertilization and embryo culture were performed, as previously described [25,34,35]. Briefly, cumulus-oocyte complexes (COC) were aspirated from follicles (3-8 mm) and identified under a stereomicroscope (10×). COC with uniform cytoplasm containing >3 layers of cumulus cells were selected for further processing. COC were washed (3×) in maturation medium (5% calf serum v/v, 0.5 µg/mL FSH, 5 µg/mL LH and 0.05 µg/mL gentamycin in TCM199). Approximately 20 COC were pipetted into a 100 µL droplet of maturation medium and incubated under mineral oil at 38.5°C and 5% CO_2_ in air, for 22 h. Frozen-thawed semen samples were centrifuged (2000× g) through Percoll density gradients (45% and 90%) in a 15 mL conical tube for 15 min and pellet was diluted to 3×10^6^ sperm/mL with Brackett-Oliphant (BO) fertilization medium [25]. Mature COC were washed (3×) with BO medium containing 10% (w/v) bovine serum albumin (BSA), placed in 100 µL droplets possessing 300,000 sperm, and incubated at 38.5°C and 5% CO_2_ in air, for 18 h (Day 0 = day of IVF). After incubation, the presumptive zygotes were denuded by repeated pipetting while washing (3×) in 100 µL droplets of Charles Rosenkrans1 + amino acids culture media (CR1aa) + 5% v/v calf serum under mineral oil at 38.5°C, 5% CO_2_, 5% O_2_ and 90% N_2_. Cleavage and blastocyst rates were evaluated on Day 2 and 8, respectively, and reported based on the number of oocytes submitted to IVF. This trial was conducted four times on different dates (replicates).

#### Statistical analysis

Values are expressed as mean±standard error of the mean (SEM) unless otherwise stated. In Experiment 1, 3 × 2 factorial analysis was used to study the effect of extenders (TEYG, animal-CCTG and plant-CCTG), freezing methods (routine and fast) and their interactions on the post-thaw sperm characteristics. In Experiments 2 and 3, factorial analysis (2 × 2) was used to determine the effect of extenders and freezing methods. If main effects or their interaction were significant (P ≤ 0.05), means were compared by Tukey’s test. *In vitro* cleavage and blastocyst rates, and *in vivo* pregnancy rates were compared among groups by binomial generalized linear mixed model analysis of variance. Data analyses were performed using the R Project (R version 3.3.1, R Foundation for Statistical Computing, Vienna, Austria).

## Results

### Experiment 1: Effect of extender, origin of cholesterol and freezing method on post-thaw sperm quality and *in vitro* fertility of beef bulls

#### Experiment 1a (Simmental bulls)

Extender (TEYG, animal-CCTG or plant-CCTG) × freezing method (routine or fast) interaction was significant on post-thaw total and progressive motility at 0 and 2 h (Table 3). The TEYG NE-FF group had the lowest post-thaw total and progressive motility compared to other treatment groups. Post thaw motility did not differ between animal- and plant-origin cholesterol.

**Table 3.** Post-thaw sperm motility of beef semen diluted in TEYG, animal-CCTG or plant-CCTG extender and frozen with routine or fast freezing method. Each value represents the mean±SEM of five pooled ejaculates (replicates), from five Simmental bulls (Experiment 1a).

| Sperm characteristics | Time | Routine method (RE-RF) |  |  | Fast method (NE-FF) |  |  |
| --- | --- | --- | --- | --- | --- | --- | --- |
|  |  | TEYG | Animal-CCTG | Plant-CCTG | TEYG | Animal-CCTG | Plant-CCTG |
| Total motility (%) | 0 | 48±5.0<sup>a</sup> | 50±4.7<sup>a</sup> | 52±5.3<sup>a</sup> | 24±3.2<sup>b</sup> | 47±3.3<sup>a</sup> | 43±3.1<sup>a</sup> |
|  | 2 | 51±2.6<sup>a</sup> | 50±4.8<sup>a</sup> | 46±7.6<sup>a</sup> | 21±2.5<sup>b</sup> | 45±4.7<sup>a</sup> | 40±8.8<sup>a</sup> |
| Progressive motility (%) | 0 | 38±3.5<sup>a</sup> | 43±4.9<sup>a</sup> | 44±5.0<sup>a</sup> | 20±3.1<sup>b</sup> | 38±2.3<sup>a</sup> | 36±2.0<sup>a</sup> |
|  | 2 | 40±3.8<sup>a</sup> | 43±4.5<sup>a</sup> | 40±6.8<sup>a</sup> | 14±2.1<sup>b</sup> | 37±4.2<sup>a</sup> | 32±7.3<sup>a</sup> |

<sup>ab</sup>Within rows, values with different superscripts are different ( $P < 0.05$ ).
Abbreviations: Teyg, tris-egg yolk-glycerol extender; CCTG, cholesterol-cyclodextrin tris-glycerol extender; RE-RF, routine equilibration-routine freezing; NE-FF, no equilibration-fast freezing.

Following *in vitro* fertilization of cattle oocytes with frozen-thawed sperm, cleavage and blastocyst rates ranged from 55% to 65% and 23% to 33%, respectively (Table 4). No effect of extender or freezing method was detected on cleavage or blastocyst rate.

**Table 4.** Cleavage and blastocyst rates following *in vitro* fertilization of bovine oocytes with beef semen diluted in TEYG, animal-CCTG or plant-CCTG extender and frozen with routine or fast freezing method. Each value represents four replicates conducted on different dates (Experiment 1a).

| IVF variables | Routine method (RE-RF) |  |  | Fast method (NE-FF) |  |  |
| --- | --- | --- | --- | --- | --- | --- |
|  | TEYG | Animal-CCTG | Plant-CCTG | TEYG | Animal-CCTG | Plant-CCTG |
| Cleavage rate* | 58/96<br>(60%) | 73/114<br>(64%) | 58/105<br>(55%) | 52/83<br>(63%) | 83/134<br>(62%) | 87/133<br>(65%) |
| Blastocyst rate* | 29/96<br>(30%) | 35/114<br>(31%) | 29/105<br>(28%) | 19/83<br>(23%) | 38/134<br>(28%) | 44/133<br>(33%) |

\*Cleavage and blastocyst rates were calculated based on the number of oocytes submitted to IVF.
Abbreviations: TEYG, tris-egg yolk-glycerol extender; CCTG, cholesterol-cyclodextrin tris-glycerol extender; RE-RF, routine equilibration-routine freezing; NE-FF, no equilibration-fast freezing.

#### Experiment 1b (Angus bulls)

Extender × freezing method interactions were significant for all post-thaw sperm characteristics at 0 and 2 h (Table 5). Once again, all post-thaw sperm characteristics were lowest in the TEYG fast (NE-FF) group than in other groups. Post-thaw sperm total and progressive motility, and sperm with intact plasma and acrosome membranes were greater (P < 0.05) in the routine (RE-RF) than the fast (NE-FF) method. Within the fast method, post-thaw sperm characteristics at 0 h were significantly greater (P < 0.05) in plant-CCTG than TEYG extender. Within the routine method, differences between TEYG and plant-CCTG, were not significant for any sperm characteristic.

**Table 5.**
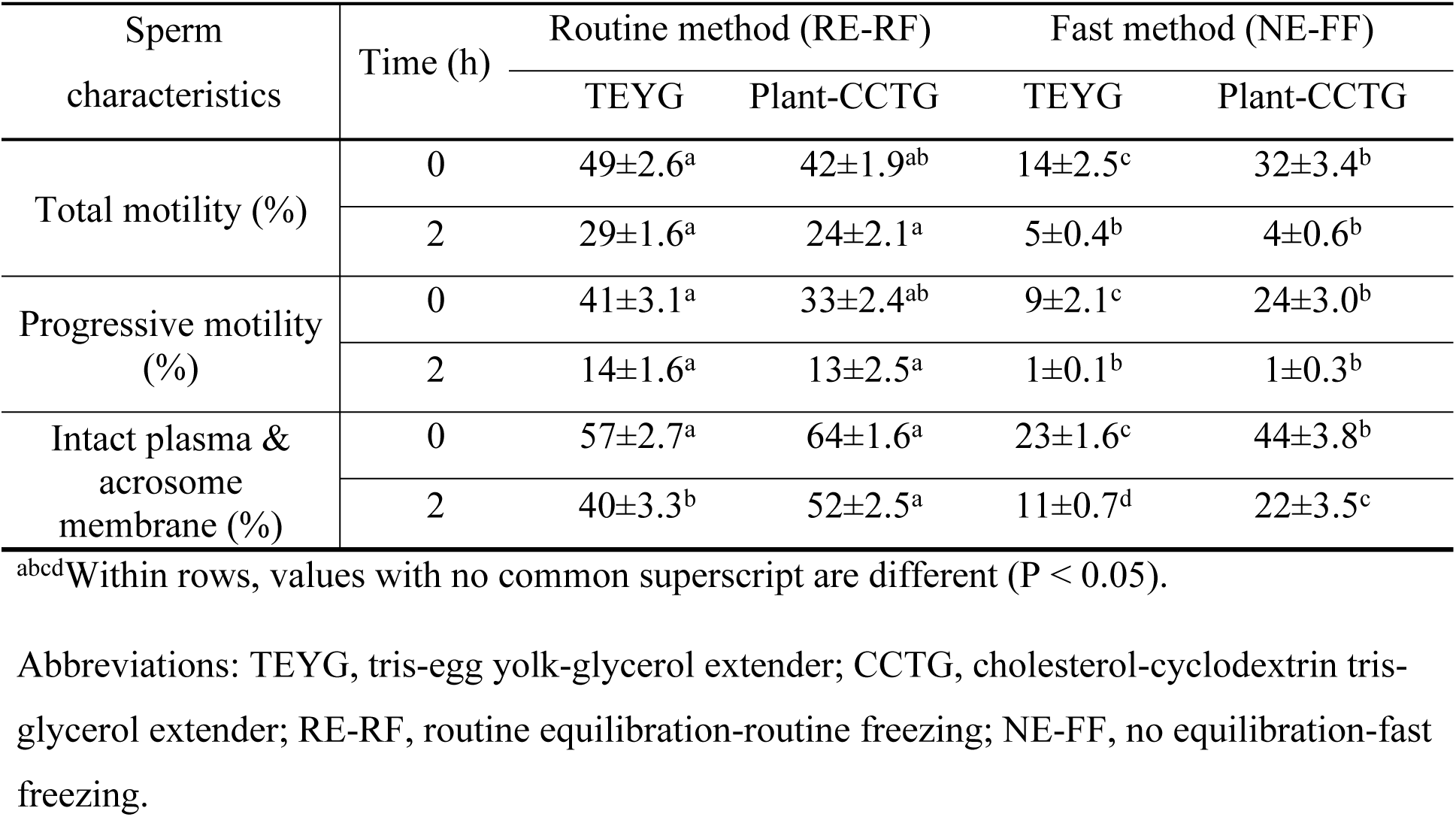
Post-thaw sperm characteristics of beef semen diluted in TEYG or plant-CCTG extender and frozen with routine or fast method. Each value represents mean±SEM of seven pooled ejaculates (replicates), from five Angus bulls (Experiment 1b).

| Sperm characteristics | Time (h) | Routine method (RE-RF) |  | Fast method (NE-FF) |  |
| --- | --- | --- | --- | --- | --- |
|  |  | TEYG | Plant-CCTG | TEYG | Plant-CCTG |
| Total motility (%) | 0 | 49±2.6<sup>a</sup> | 42±1.9<sup>ab</sup> | 14±2.5<sup>c</sup> | 32±3.4<sup>b</sup> |
| | 2 | 29±1.6<sup>a</sup> | 24±2.1<sup>a</sup> | 5±0.4<sup>b</sup> | 4±0.6<sup>b</sup> |
| Progressive motility (%) | 0 | 41±3.1<sup>a</sup> | 33±2.4<sup>ab</sup> | 9±2.1<sup>c</sup> | 24±3.0<sup>b</sup> |
| | 2 | 14±1.6<sup>a</sup> | 13±2.5<sup>a</sup> | 1±0.1<sup>b</sup> | 1±0.3<sup>b</sup> |
| Intact plasma & acrosome membrane (%) | 0 | 57±2.7<sup>a</sup> | 64±1.6<sup>a</sup> | 23±1.6<sup>c</sup> | 44±3.8<sup>b</sup> |
| | 2 | 40±3.3<sup>b</sup> | 52±2.5<sup>a</sup> | 11±0.7<sup>d</sup> | 22±3.5<sup>c</sup> |

<sup>abcd</sup> Within rows, values with no common superscript are different ( $P < 0.05$ ).
Abbreviations: TEYG, tris-egg yolk-glycerol extender; CCTG, cholesterol-cyclodextrin tris-glycerol extender; RE-RF, routine equilibration-routine freezing; NE-FF, no equilibration-fast freezing.

#### Experiment 2: Effect of extender and freezing method on post-thaw sperm quality of bison bulls

Bison sperm in the TEYG NE-FF group also had the lowest (P < 0.05) total and progressive motility, and sperm with intact plasma and acrosome membrane integrity compared to other treatment groups at 0 h post-thaw (Table 6). There was no difference in post-thaw sperm characteristics between TEYG and plant-CCTG in the routine group (RE-RF), at 0 and 2 h. Sperm extended in TEYG or plant-CCTG extenders, and underwent the fast method (NE-FF) yielded lower sperm motility and sperm with intact plasma membrane integrity and normal acrosomes at 2 h, compared to routine group (RE-RF, P < 0.05).

**Table 6.** Post-thaw sperm characteristics of bison semen diluted in TEYG or plant-CCTG extender and frozen with a routine or fast method. Each value represents mean±SEM of five pooled ejaculates (replicates), from five bison bulls (Experiment 2).

| Sperm characteristics | Time (h) | Routine method (RE-RF) |  | Fast method (NE-FF) |  |
| --- | --- | --- | --- | --- | --- |
|  |  | TEYG | Plant-CCTG | TEYG | Plant-CCTG |
| Total motility (%) | 0 | 45±1.0<sup>a</sup> | 51±4.1<sup>a</sup> | 10±1.2<sup>b</sup> | 39±4.8<sup>a</sup> |
| | 2 | 32±2.8<sup>a</sup> | 31±2.1<sup>a</sup> | 3±1.0<sup>b</sup> | 3±0.4<sup>b</sup> |
| Progressive motility (%) | 0 | 37±1.0<sup>a</sup> | 45±4.4<sup>a</sup> | 5±1.0<sup>b</sup> | 33±4.7<sup>a</sup> |
| | 2 | 26±2.0<sup>a</sup> | 24±2.4<sup>a</sup> | 1±0.2<sup>b</sup> | 1±0.4<sup>b</sup> |
| Intact plasma membrane & acrosome (%) | 0 | 47±1.5<sup>a</sup> | 52±6.1<sup>a</sup> | 17±1.9<sup>c</sup> | 30±4.6<sup>b</sup> |
| | 2 | 39±1.9<sup>a</sup> | 42±4.7<sup>a</sup> | 15±1.7<sup>b</sup> | 15±2.8<sup>b</sup> |

<sup>abc</sup>Within rows, values with no common superscript are different ( $P < 0.05$ ).
Abbreviations: TEYG, tris-egg yolk-glycerol extender; CCTG, cholesterol-cyclodextrin tris-glycerol extender; RE-RF, routine equilibration-routine freezing; NE-FF, no equilibration-fast freezing.

#### Experiment 3: Effect of extender and modified freezing method on post-thaw sperm quality and *in vivo* fertility of beef bulls

There was no effect of extender or interaction between main effects on post-thaw sperm characteristics. However, sperm motility, and membrane and acrosome integrity were lower in the modified than routine freezing method (P < 0.05; Table 7).

**Table 7.** Post-thaw sperm motility, and plasma membrane integrity and normal acrosomes of beef semen diluted in TEYG or plant-CCTG and frozen using a routine or modified method. Each value represents the mean±SEM of ten pooled ejaculates (replicates) from six Angus bulls (Experiment 3).

| Sperm characteristics | Time (h) | Routine method (RE-RF) |  | Modified method (NE-RF) |  |
| --- | --- | --- | --- | --- | --- |
|  |  | TEYG | Plant-CCTG | TEYG | Plant-CCTG |
| Total motility (%) | 0 | 45±3.4<sup>a</sup> | 43±1.6<sup>a</sup> | 37±3.4<sup>b</sup> | 37±3.4<sup>b</sup> |
| | 2 | 33±3.8<sup>a</sup> | 31±2.0<sup>a</sup> | 25±4.0<sup>b</sup> | 22±3.2<sup>b</sup> |
| Progressive motility (%) | 0 | 35±3.6<sup>a</sup> | 34±2.2<sup>a</sup> | 26±3.5<sup>b</sup> | 27±3.5<sup>b</sup> |
| | 2 | 23±3.9<sup>a</sup> | 21±2.2<sup>a</sup> | 16±3.1<sup>b</sup> | 12±3.4<sup>b</sup> |
| Intact membranes (%) | 0 | 68±2.4<sup>a</sup> | 64±3.4<sup>a</sup> | 56±8.6<sup>b</sup> | 56±3.9<sup>b</sup> |
| | 2 | 56±3.6<sup>a</sup> | 52±4.4<sup>a</sup> | 38±9.5<sup>b</sup> | 37±4.8<sup>b</sup> |

<sup>ab</sup>Within rows, values with different superscripts are different ( $P < 0.05$ ).
Abbreviations: TEYG, tris-egg yolk-glycerol extender; CCTG, cholesterol-cyclodextrin tris-glycerol extender; RE-RF, routine equilibration-routine freezing; NE-RF, no equilibration-routine freezing.

In Fertility trial 1, no difference was detected between CIDR and PRID-Delta groups in ovulation rate (102/105 [97%] vs 101/101 [100%] or interval to ovulation from GnRH treatment (30.8±1.5 h vs 32.3±1.6 h). Pregnancy rate was greater with PRID-Delta (74/101 [73%] compared to CIDR (63/105 [60%], P ≤ 0.05), and TEYG routine (RE-RF; 78/104 [75%]) compared to plant-CCTG modified (NE-RF; 59/102 [58%]; Table 8).

**Table 8.**
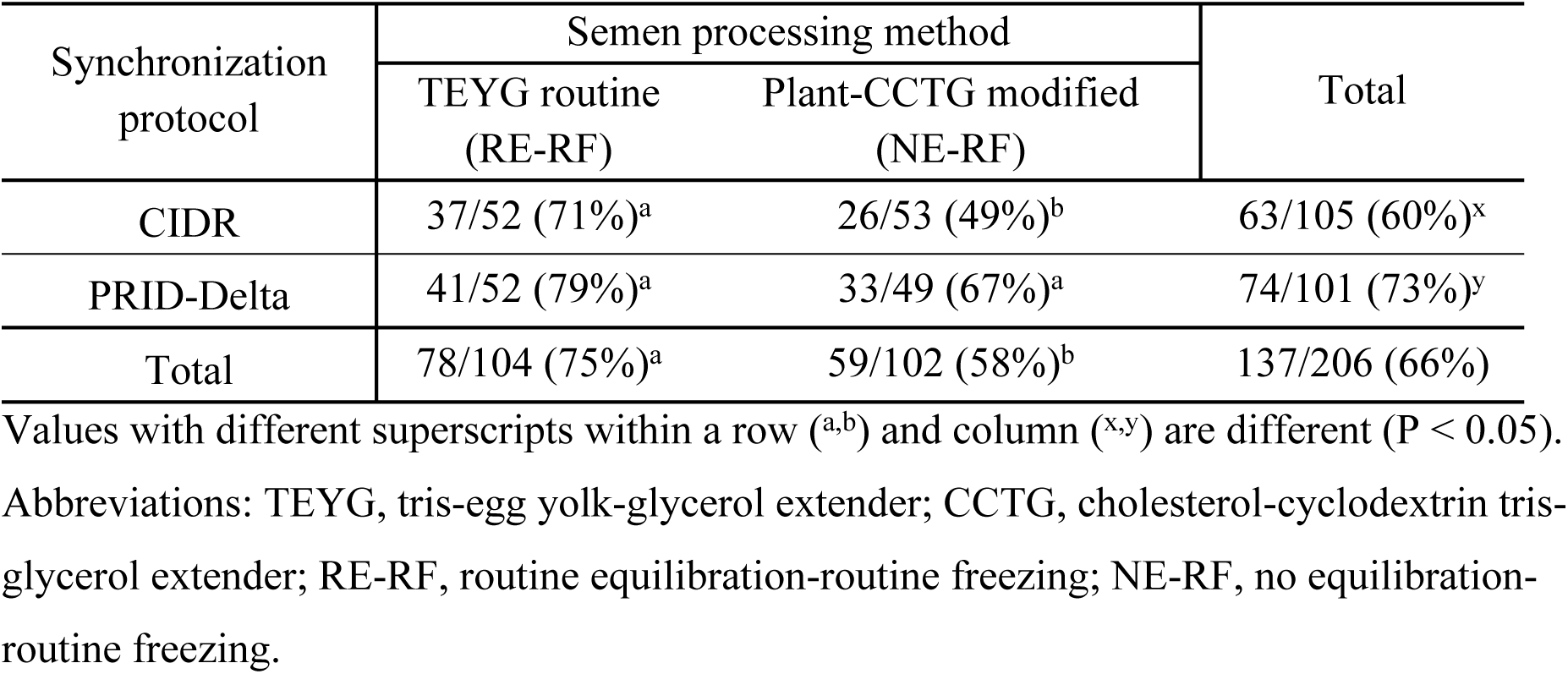
Pregnancy rate (number pregnant/number inseminated) in beef cattle following fixed-time artificial insemination using CIDR vs PRID-Delta as a source of progesterone for ovarian synchronization and using either a routine (TEYG, RE-RF) or modified (plant-CCTG, NE-RF) semen processing method (Fertility trial 1).

In Fertility trial 2, the overall ovulation rate did not differ between the 60 h and 72 h groups (64/65 [98%] vs 40/41 [98%], respectively), but a greater proportion of cows had ovulated at the time of insemination in the 72 h group than in the 60 h group (16/41 [39%] vs 6/65 [9%]; P ≤ 0.05). A 2 × 2 interaction effect of insemination timing and semen group on pregnancy rate was significant (P < 0.05; Table 9). Plant-CCTG modified (NE-RF) with 60 h insemination timing had the lowest pregnancy rate compared to other groups (P ≤ 0.05). There was no difference in pregnancy rate between semen extender groups when insemination was done at 72 h.

**Table 9.**
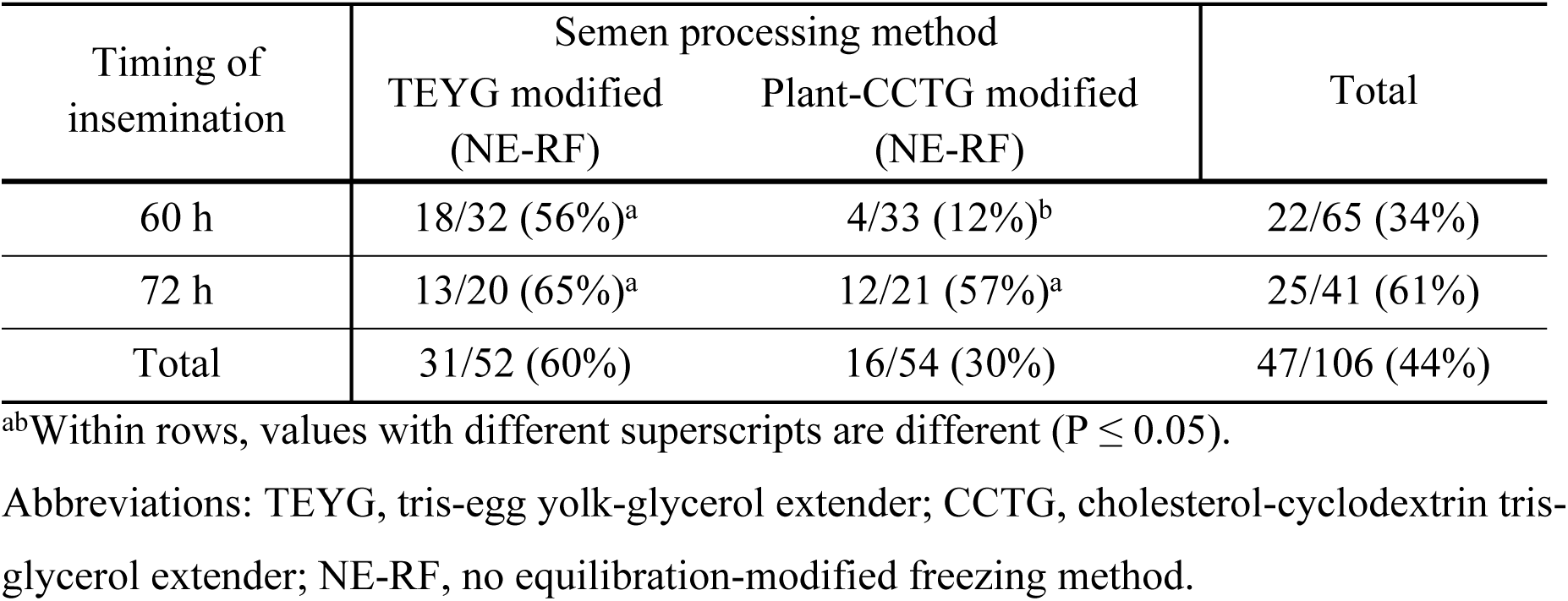
Pregnancy rate (number pregnant/number inseminated) in beef cattle following fixed-time artificial insemination with modified semen (NE-RF) in TEYG or plant-CCTG extenders at 60 h vs 72 h after progesterone device removal (Fertility trial 2).

## Discussion

In the present study, the effects of semen processing during cryopreservation were examined by altering equilibration time and freezing rate in different semen extenders. Results demonstrated that beef and bison semen can be frozen without equilibration in a cholesterol-based (animal product-free) extender using a fast freezing method (no equilibration-fast freezing, NE-FF) in 14 min, with acceptable post-thaw sperm motility, plasma membrane integrity, normal acrosomes and *in vitro* blastocyst formation. In contrast, semen diluted in TEYG extender, directly cooled to 4°C without equilibration and frozen with a fast freezing method (NE-FF) yielded the lowest post-thaw semen quality. Surprisingly, semen diluted in TEYG extender and frozen with a modified method (no equilibration-routine freezing, NE-RF) yielded comparable post-thaw sperm motility and *in vivo* fertility (up to 65%).

There is no clear consensus in the literature on equilibration time for bull semen. The extended semen is routinely equilibrated for 90-min at 4°C to allow the low density lipoproteins in egg yolk to interact with sperm plasma membranes [36]. These lipoproteins protect sperm from cold shock and lipid-phase transitions that occurs around 15°C during cooling to 4°C [2,16], and replenish phospholipids lost during cryopreservation [37]. In contrast, bull and bison semen diluted in animal- or plant-CCTG was successfully frozen with or without a 90-min equilibration period in the current study. Earlier, we reported that exogenous cholesterol (plant-origin) extender can be used to cryopreserve cattle and bison semen [11,25]. Cholesterol increases the fluidity of sperm plasma membranes and minimizes membrane lipid phase transitions during initial cooling [38]. Data from Experiment 1 showed no difference in post-thaw sperm motilities between animal- and plant-origin cholesterol as expected, since they have identical structural formulas. Therefore, plant cholesterol was used in subsequent experiments as it represents a biosecure alternative to cholesterol from sheep’s wool.

Damage to sperm during cryopreservation is twofold. Above 0°C, sperm plasma membrane undergoes phase transition around 15°C. Below 0°C, intracellular ice formation damages subcellular structures [39]. In the routine (RE-RF) method, the diluted semen was equilibrated at 4°C for at least 90 min and frozen as described in previous studies [11,25,30]; whereas in the fast and modified methods, semen was cooled @ -1°C/min from 22°C to 10°C or 4°C in a programmable cell freezer eliminating routinely used 90-min equilibration. In addition, the freezing rate was changed from routine (double-ramp: @ -3°C/min from 4°C to -10°C and @ -40°C/min from -10°C to -80°C) to fast (single ramp: @ -40°C/min from 10°C to -80°C) freezing. In Experiments 1 and 2 on beef and bison semen respectively, the TEYG fast-freezing (NE-FF) group had the lowest post-thaw sperm quality (motility, plasma membrane integrity and normal acrosomes) compared to other groups (TEYG-routine, plant-CCTG-routine and CCTG-fast). In Experiment 3, the cryopreservation process was modified such that equilibration was omitted, but freezing rate below 0°C remained as per routine (NE-RF). Post-thaw quality of semen in TEYG modified and plant-CCTG modified (NE-RF) groups were similar. Previous work revealed that cooling rate (0.1 or 4.2°C/min) had no effect on post-thaw sperm characteristics or *in vivo* fertility using egg yolk-based extender in dairy bulls [40]. Therefore, cryodamage associated with fast method was likely due to ice nucleation [26] occurring around - 40°C in buffalo bull semen [41]. The freezing rate (-3°C/min; ramp 1) used in routine and modified groups in the present study allowed more gradual cell dehydration and ice nucleation [42]. In Experiments 1 and 2, the freezing rate of -40°C/min in ramp 1 was >10× faster, leading to inadequate cell dehydration and subsequent ice nucleation of intracellular free water and crystal formation. Interestingly, CCTG extender provided protection against fast freezing, presumably by adding exogenous cholesterol to sperm plasma membrane, though the exact protective mechanism of exogenous cholesterol in bull and bison sperm is yet to be determined. In previous reports, post-thaw sperm quality and *in vivo* fertility varied with the use of different equilibration times (24 to 90 h) at 4°C [16,18,19,43]. Contrary to results of the present study, others found that 0 h equilibration of bull semen yielded low sperm motility compared to 4 h [43]. We anticipated that the equilibration process of semen diluted in egg yolk extender is slow, necessitating a prolonged equilibration time [18]. However, our results demonstrate that the long equilibration time with egg yolk extender is required for successful cryopreservation. *In vitro* cleavage and blastocyst rates were unaffected due to extender or freezing method in the present study. Prior to *in vitro* fertilization of cattle oocytes, frozen-thawed semen is routinely washed through a density gradient to select for viable sperm, and equal number of viable sperm are co-incubated with *in vitro* matured oocytes [25,34,35]. The lack of difference in cleavage and blastocyst rates may have been a result of high number of viable sperm per fertilization droplet that compensates for fertility-associated sperm defects [44–46]. The compensatory effect of sperm number was minimized when sperm number per fertilization droplet was reduced from 300,000 to 30,000, and a difference in blastocyte rate due to treatment was more conspicuous [47]. A conventional *in vitro* fertility trial may not be a reliable assay of *in vivo* fertility [25], until the sperm concentration per droplet is reduced.

North American bison are a significant wildlife reservoir for zoonotic diseases such as bovine tuberculosis and brucellosis. Concerted efforts are underway to conserve and redistribute disease-free bison genetics among long-isolated populations. To this end, it is important to produce biosecure frozen semen to prevent the spread of disease, and to be able to optimize a cryopreservation process that is applicable in field conditions. Fast freezing comes at a cost of increasing sperm damage, particularly with the use of TEYG extender and with the more sensitive bison semen. Like beef bulls, post-thaw sperm characteristics of bison semen diluted in TEYG extender and frozen with the fast method (NE-FF) were adversely affected compared to other treatment groups. However, overall post-thaw sperm quality characteristics were lower in bison compared to beef bulls. This is in consistent with previous findings that bison sperm are more prone to membrane damage compared to cattle [25]. The rapid technique developed herein may be useful for cryopreservation of bison semen under wild conditions for *in vitro* fertilization later.

Fertility trial 1 revealed that the TEYG routine group had a higher pregnancy rate than the plant-CCTG modified group; however, in Fertility trial 2 pregnancy rates were the same between TEYG and plant-CCTG modified freezing groups when insemination was done at 72 h after progesterone device removal. The lower pregnancy rate using plant-CCTG modified method (NE-RF) inseminated at 60 h may have been a result of asynchrony between sperm viability/longevity and the timing of ovulation. Cholesterol efflux from the plasma membrane is crucial for sperm capacitation and fertilization [13,48], and oversaturation of the sperm plasma membrane with exogenous cholesterol reduced fertility potential and thus demands longer capacitation time [49]. The pregnancy rate was higher in cows synchronized with PRID-Delta than with CIDR, consistent with previous findings [50], perhaps as a result of higher circulating concentrations of progesterone with PRID than CIDR [51]. To our knowledge, this is the first report documenting optimum field fertility using semen frozen in plant-CCTG (egg yolk-free) extender without equilibration. Further investigations are required by including more cows in the fertility trial. However, 58% *in vivo* fertility (with matching ovarian synchrony) using a biosecure animal product-free extender and short processing time is comparable to cattle industry standards (50-60%), following fixed-time artificial insemination.

## Conclusion

Cattle and bison semen can be cryopreserved without equilibration at 4°C using plant-CCTG extender to achieve acceptable post-thaw sperm quality and *in vivo* fertility. Cholesterol in plant-CCTG extender requires less time than egg yolk to provide its cryoprotective effect. However, more time may be required between insemination and ovulation to maximize pregnancy rate.

Results support the stated hypothesis that a cholesterol-based semen extender obviates the need of extended equilibration for semen cryopreservation, and serves the purpose of enhanced biosecurity, particularly for beef and bison semen collection and processing under field conditions. It is important to consider the synchronization protocol and timing of artificial insemination to achieve optimum fertility in a fixed-time artificial insemination program. Additional studies are required to refine the freezing method using plant-CCTG extender supplemented with antioxidants and membrane stabilizers to improve post-thaw sperm longevity.

## Acknowledgements

Research was supported by grants from the Agriculture and Agri-Food Canada (Research Grant # AGR-14227), the Natural Sciences and Engineering Research Council of Canada, and the Bison Integrated Genomics (BIG) Project from Genome Canada’s Genomic Applications Partnership Program (GAPP) in partnership with Parks Canada and Agriculture and Agri-Food Canada (Grant # 534839). Student salary support was provided by a grant from MITACS (Grant number: IT31476). We thank Kylie Hutt, Laurence de Araujo, Lianne Price, David Moore, Justin Pifko and Jessie Hellquist for their technical assistance and the staff of the Livestock and Forage Centre of Excellence for animal management and care during the study.

